# Computing Skin Cutaneous Melanoma Outcome from the HLA-alleles and Clinical Characteristics

**DOI:** 10.1101/850677

**Authors:** Anjali Dhall, Sumeet Patiyal, Harpreet Kaur, Sherry Bhalla, Chakit Arora, G.P.S. Raghava

## Abstract

Human Leukocyte Antigen (HLA) is an essential component of the immune system which stimulates immune cells to provide protection and defense against cancer. More than thousands of HLA alleles have been reported in the literature; but, only a specific set of HLA alleles expressed in an individual. Recognition of cancer-associated mutations by the immune system depends on the presence of a particular set of alleles, that elicit an immune response to fight against cancer. It indicates that the occurrence of specific HLA alleles also affects the outcome of the cancer patients. In the current study, prediction models have been developed using 415 skin cutaneous melanoma (SKCM) patients for predicting the overall survival of patients from their HLA-alleles. It has been observed that, the presence of certain superalleles in the patients, is responsible for improved overall survival which were referred as favourable superalleles like HLA-B*55 (HR=0.15, 95% CI 0.034 to 0.67), HLA-A*01 (HR=0.5, 95% CI 0.3 to 0.8). In contrast, presence of certain superalleles in the patients is responsible for their poor survival, those superalleles were referred as unfavourable superalleles such as HLA-B*50 (HR=2.76, 95% CI 1.284 to 5.941), HLA-DRB1*12 (HR=3.44, 95% CI 1.64 to 7.2). We developed prediction models using 14 HLA-superalleles and five clinical characteristics for predicting high-risk SKCM patients and achieve HR=4.52 (95% CI 3.088-6.609) with p-value = 8.01E-15. Lastly, we provide a web-based service to community for predicting the risk in SKCM patients (https://webs.iiitd.edu.in/raghava/skcmhrp/)

## 1 Introduction

HLA complex is highly polymorphic genetic region, located on chromosome 6, precisely 6p21.3 region (1,2). Major histocompatibility complex (MHC) proteins encode more than 200 immune-related genes, from which, approximately 40 genes were associated with the development of leukocyte antigens, *i.e*. Class I and Class II HLA genes (3). Out of which, class I genes encode proteins which present antigens (intracellular peptides) to CD8^+^ T lymphocytes, while, class II genes encode proteins which are present on antigen-presenting cells (APC) that regulate the proliferation and initiation of CD4^+^ T cells(4,5). Furthermore, Class I HLA genes are of three types, *i.e.* A, B and C, while class II HLA genes are of five types, which include DR, DP, DM, DQ, and DO. Class I complex generally located on the nucleated cell surface, and Class II genes expressed on the specific cells such as monocytes, macrophages, and dendritic cells also known as APCs, B lymphocytes and activated T cells (2).

Human Leukocyte Antigen (HLA) molecules play a major/significant role in the induction and regulation of immune responses. The role of HLA class I molecules has been implied in tumor resistance to apoptosis (6). Moreover, recent findings suggest that the altered expression of HLA molecules was associated with metastatic progression and poor prognosis in tumor (7–9). The modification of surface molecules, lack of co-stimulatory molecules, production of immunosuppressive cytokines, and alterations in HLA molecules are some primary escape mechanisms used by tumor cells to evade the immune response(10), which can directly affect the survival of an individual. Figure 1 represents how the survival of the patients can get affected if HLAs fails to recognize the tumor cells, which is ultimately responsible for the activation of the immune system. Previous studies reveal that skin cutaneous melanoma has been reported to be the most threatening and fatal form of skin cancer and scrutinized multi-omics signatures for the progression of this malignancy (11–13). It has been shown that if melanoma is detected at an early stage, the overall survival rate is 95%; but, once it is metastasized (lesion thickness >4mm); they are tough to cure and the survival rate is reduced to less than 50% (14,15). Melanoma tumor cells escape the immune checkpoints and proliferate at a higher rate than normal tissue cells (16). Further, it is categorized as an immunogenic tumor as it’s lesions have been found to have signatures of several immune escape mechanisms such as downregulated expression of HLA molecules, secretion of cytokines like IL10 and loss of tumor-specific antigens (17).

**Figure 1:**
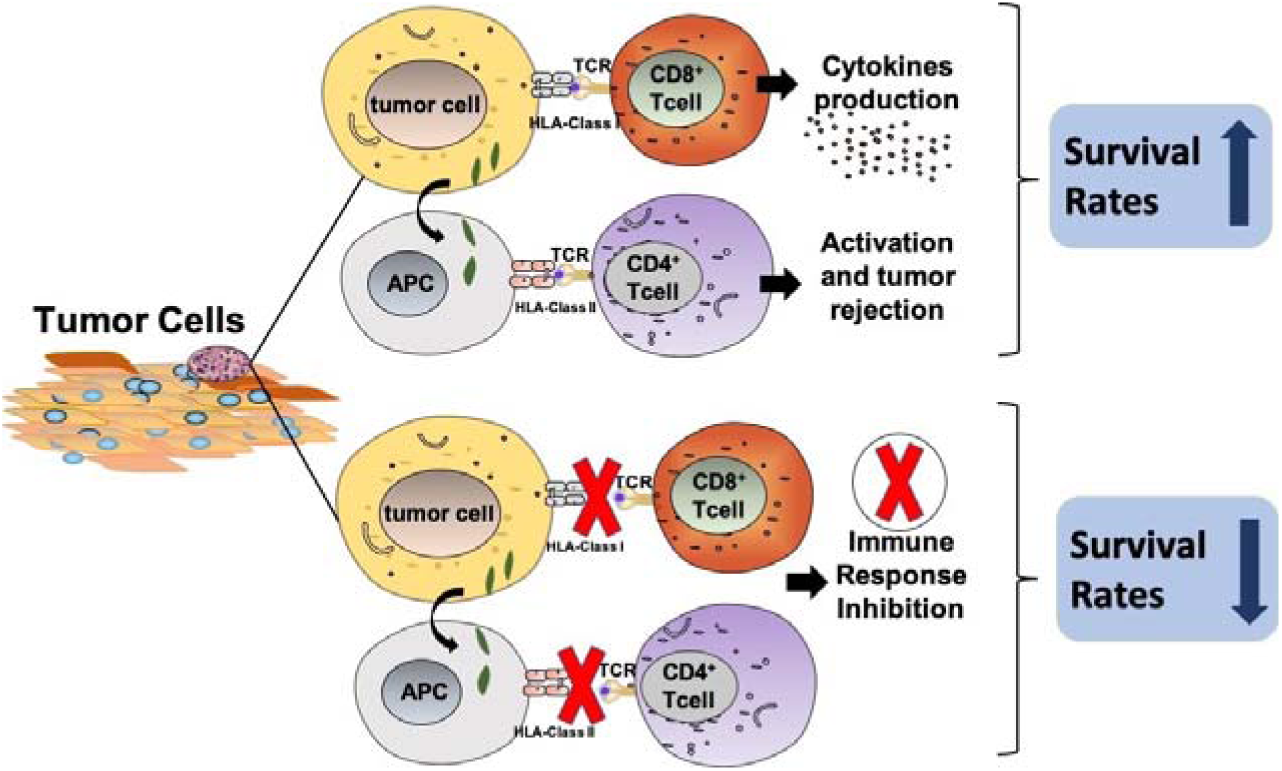
The identification of tumor cells by CD8^+^ cytotoxic T cells and CD4^+^ T helper cells via HLA class I and II molecules, respectively.

For instance, the downregulation of class I antigens was associated with poor prognosis and inadequate treatment in melanoma cases (18–20). Moreover, recent studies demonstrate the importance of HLA alleles in the prognosis of melanoma, such as the loss of heterozygosity of HLA class I alleles (HLA-B*15:01) was shown to be related with poor survival outcome. Besides, HLA-C alleles and HLA-B44 supertype were shown to enhance the overall survival (21–23), thereby claiming that these molecules could be considered as prognostic markers for melanoma. Thus, it is vital to analyse the role of class I and II antigens in the survival of melanoma patients. With the knowledge of accurate HLA genotyping, one can design immunotherapy-based prognostic biomarkers and personalized vaccines against cancer.

In the current study, we have made an attempt to understand the role of HLA (Class I and II) alleles and superalleles in the survival of the skin cutaneous melanoma (SKCM) patients using TCGA-SKCM’s cohort. Here, firstly we have performed HLA-genotyping of patients for the Class I and II alleles, followed by their assignment to the superalleles groups. Subsequently, we categorized the superalleles into survival favourable and unfavourable superallele groups based on the impact of their presence on the survival of the patients. Further, we have developed survival prediction models employing key superalleles and clinical features of the patients by using different machine learning techniques. Eventually, to serve the scientific community, we have developed a webserver for the prediction of low-risk and high-risk patients’ groups based on the HLA-Superalleles and Clinical features.

## 2 Methods and Materials

### 2.1 Study Design and Dataset Collection

The workflow of our study is illustrated in Figure 2. The description of each step given below.

**Figure 2:**
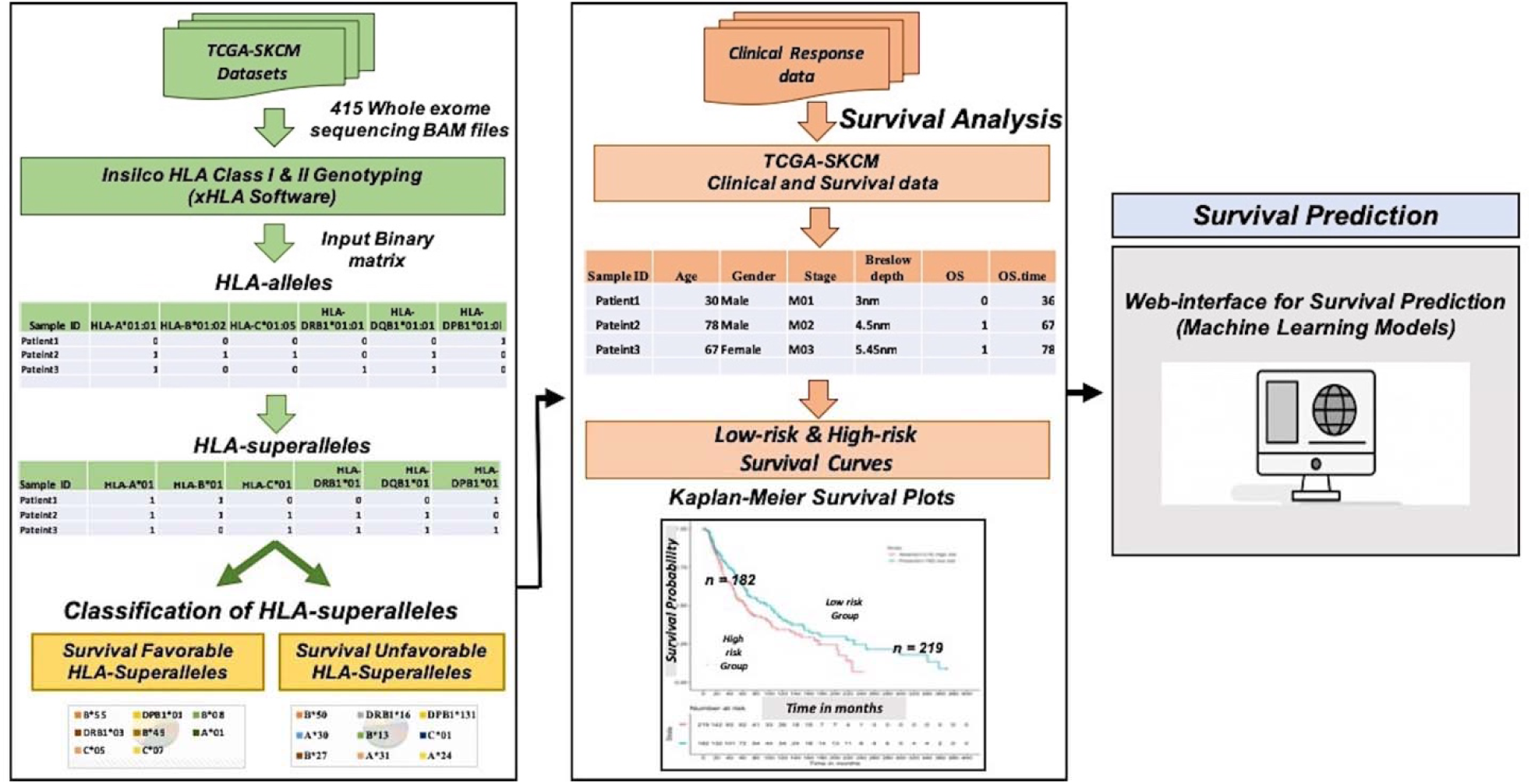
Work flow present overall architecture of this study

### 2.2 Skin cutaneous melanoma patient’s data

We have downloaded the SKCM controlled access dataset from GDC data portal. Specifically, the whole-exome sequencing (WXS) BAM files of individual melanoma patient were downloaded (under the approval of dbGap (Project No. 17674)) according to the Genome Data Commons protocols (24) with the help of in-house HPC facility and scripts. Clinical information for 470 SKCM patients also obtained, that includes age, gender, stage, tumor status, treatment status, Breslow Depth, vital status, Overall Survival (OS) time, etc. using TCGA assembler 2 (25,26). We were able to extract the HLA-typing information for 415 patients out of 470 TCGA-SKCM samples only. Out of 415 samples, fourteen SKCM samples lack overall survival information. In summary, we used 401 SKCM-patients for which complete survival information is available with exome sequencing data.

### 2.3 HLA Typing and Assignment into Superalleles

After downloading the whole exome BAM files of SKCM-patients from TCGA, chromosome 6 was extracted from these BAM files using SAMtools package (27). In next step, we used xHLA software (https://github.com/humanlongevity/HLA) for HLA genotyping from chromosome 6. In this study, four-digit HLA typing was performed for each patient for the assignment of both Class I (-A, -B, -C) and Class II (-DP, -DQ, -DR) HLA genes. Further, an allele is assigned to HLA-superallele on the basis of common family alleles (Field F2), i.e. HLA-alleles were grouped to HLA-superalleles on the basis of similar HLA-Gene (-A, -B, -C, -DPB1, -DQB1, -DRB1) and Field1 (F1) (which represents the allele of a particular gene)(28), the complete representation is given in Figure 3 and Supplementary Table S2.

**Figure 3:**
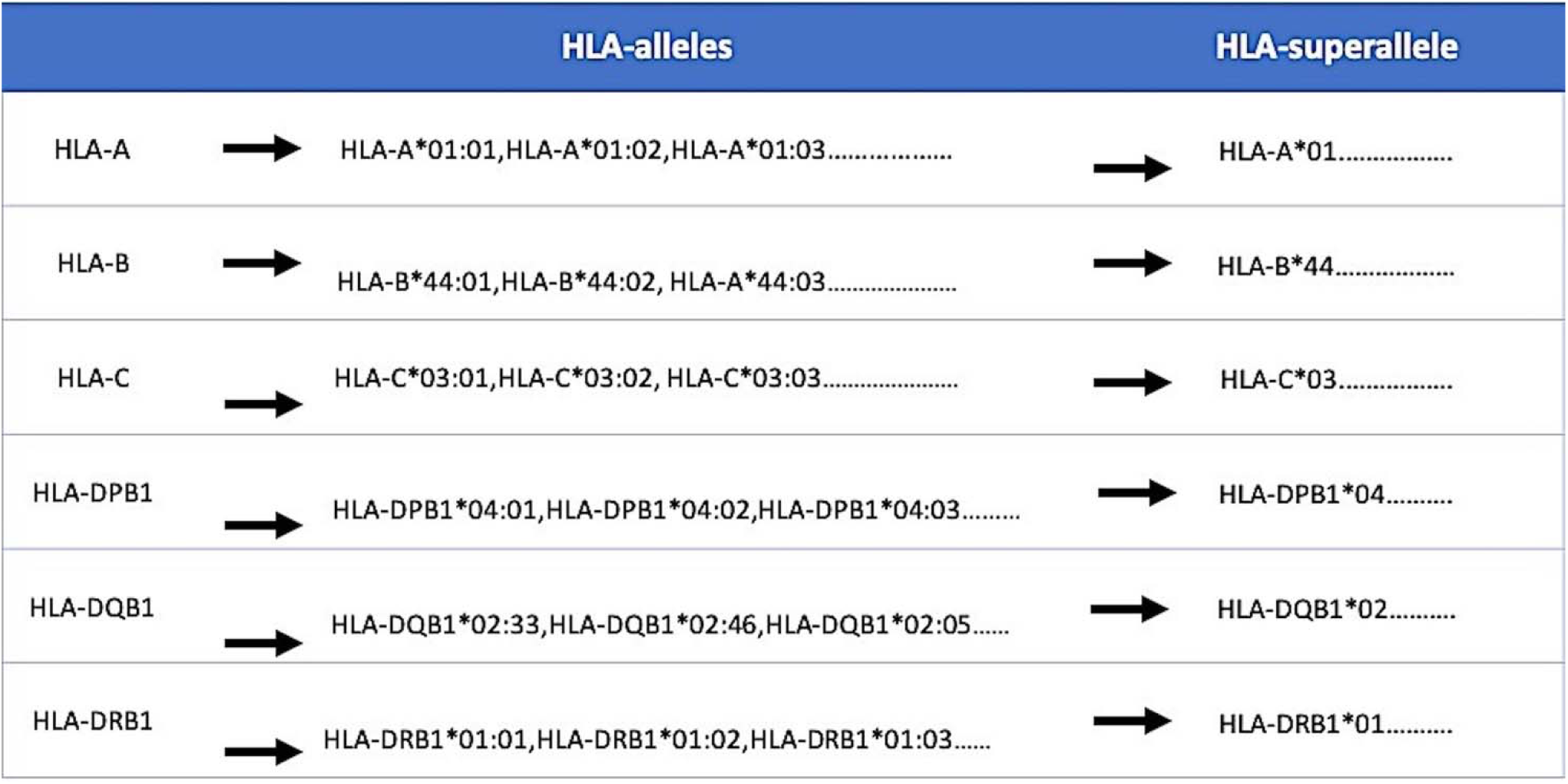
Representation of HLA-superalleles on the basis of common HLA-gene (-A, -B, -C, -DPB1, -DQB1, -DRB1) and Field 1 (F1). F1 and F2 represent the allele group and specific HLA-protein respectively.

### 2.4 Categorization of HLA-Superalleles

Here, we categorized all HLA alleles into favourable and unfavourable groups based on the impact of their presence on the survival of patients, i.e. whether the presence of superalleles either improve or decrease the survival. Towards this, firstly, all patients were divided into two groups, i.e. patients having a particular allele and the patients lacking it; subsequently, the mean survival of patients was computed in each group. Further, an allele is assigned as survival favourable allele if the mean survival of the patients having this allele is more than the mean survival of patients lacking this allele. Similarly, an allele is assigned as an unfavourable allele, if the mean survival of the patients containing this allele is lower than the mean survival of patients without this allele. It has been observed that an individual allele is only present in the limited number of patients; thus, grouping based on the occurrence of allele will be skewed. Therefore, eventually, we analyse the presence and absence of HLA-superalleles in the patients and assigned them in favourable (SF) and unfavourable (SU) superalleles groups. Notably, we considered only those superalleles, that must be present in atleast ten samples before assigning it into any of these groups. Further, to study the overall impact of the presence of SF and SU superalleles, we combine SF and SU superalleles and prepare a matrix; where, we assign a score +1 if unfavourable and -1 if favourable superallele is present in an SKCM patient, otherwise 0. Eventually, all the scores are cumulatively added to generate a single score called “Risk Score”. Subsequently, threshold-based methods have been developed using these superalleles as features. Finally, we assign a patient on high-risk if the score is more than threshold of Risk Score, otherwise low-risk.

### 2.5 Survival Analysis

In the current study, “Univariate” and “Multivariate” survival analysis is performed by using Cox Proportional Hazard (Cox PH) models implementing ‘survival’ package in R (V.3.5.1). To understand the impact of each variables like age, tumor stage, tumor status, sex, class I, II HLA-alleles, HLA-superalleles and Risk Score in the prognosis of SKCM patients, univariate analysis is performed. Further, to determine the combined effect of multiple factors such as age, tumor stage, tumor status, sex and class I, II HLA-superalleles, multivariate survival analysis is performed. The log-rank test was used for the estimation of significant survival distributions between high-risk and low-risk groups in terms of p-value. To demonstrate the performance of models graphically, high-risk and low-risk groups are represented by Kaplan-Meier plots (29).

### 2.6 Development of Prediction Models

#### 2.6.1 Models based on machine learning techniques

In the current study, various machine learning techniques have been implemented to develop regression models for the survival prediction in melanoma patients. These machine learning techniques include Random Forest (RF) (30), Ridge, Lasso (31), and Decision tree (DT) (32). Most of these techniques were implemented using python-library scikit-learn (33). To develop prediction models, we used a wide range of features that include HLA-superalleles, clinical characteristics of the patients like age, gender, stage, tumor status, Breslow depth, and combination of both.

#### 2.6.2 Wrapper based feature selection method

Here, a recursive feature selection model was developed by adding one-by-one HLA-superalleles to the clinical features based on the performance of each model. Then, survival time was predicted and followed by computation of Hazard Ratio (HR) for each combination. Briefly, every time input matrix was updated by adding a new column having HLA-superallele, which had the HR just higher than that of the previous input matrix. We repeat this process until there is no further improvement in HR. Finally, we are left with the matrix which attained the highest HR. Subsequently, this matrix was used to build the final prediction model for estimation of OS time.

### 2.7 Evaluation of models

#### 2.7.1 Five-fold cross-validation

In order to avoid the over optimisation in the training of models, we used standard five-fold cross-validation (34). In brief, all instances are randomly divided into five sets; where, four sets are used for the training and remaining fifth set for testing. This process is repeated five times so that each set is used for testing atleast once. The final performance is calculated by averaging the performance on all five sets.

#### 2.7.2 Parameters for measuring performance

The major challenge in these types of studies is to use appropriate parameters for evaluating the performance of models. In this study, we used standard parameter Hazard Ratio (HR) for measuring the performance of the models. HR is a measure of the effect of an intervention on an outcome of interest over time. Our regression models segregate patients into high-risk and low-risk groups by taking median cut-off. In order to evaluate our model, we compute HR from the predicted group of patients (high-risk or low-risk patients). Besides, we also measure the confidence interval (CI) with HR and reported HR at 95% CI. In order to measure the significance of prediction, we also calculate p-value by using log-rank test. These parameters were implemented previously in various similar kind of studies (35,36).

## 3 Results

### 3.1 Distribution of HLA alleles

We have extracted 367 HLA alleles for 415 SKCM-patients from the HLA-genotyping of SKCM cohort using xHLA software (37). Out of these 367 alleles, 237 belong to HLA-Class I genes (-A,-B,-C) and 130 alleles correspond to Class II genes (-DP, -DQ, -DR). We compute the frequency distribution of different alleles in the patients. Due to heterogeneity in HLA-genes, all alleles are not found in an individual, so the frequency of alleles vary in each patient (38). As shown in Figure 4A, out of 415 patients only 357 patients have all six alleles, 45 patients have five alleles of HLA Class I (-A, -B, -C) genes. Most of the patients have all six class I alleles, only few patients (around 13) have less than three alleles. In case of HLA Class II genes (-DP, -DQ, -DR), only 264 patients have all six alleles. Most of the alleles present only in a single patient; 134 in case of Class I and 61 in case of Class II. Only four alleles of Class I are present in more than 100 patients. Similarly, in case of Class II, only 5 alleles are present in more than 100 patients as shown in Figure 4B and 4C, respectively. The complete frequency distribution of class I and class II alleles in the SKCM-patients is given in Supplementary Table S3. Among them, the most abundant (present in >= 20% population) class-I and class-II HLA alleles include HLA-A*02:01, HLA-A*01:01, HLA-C*07:02, HLA-C*07:01, HLA-B*07:02, HLA-A*03:01, HLA-DPB1*04:01,HLA-DQB1*03:01, HLA-DQB1*02:01, HLA-DPB1*02:01, HLA-DRB1*07:01, HLA-DRB1*05:01, HLA-DRB1*15:01, respectively, as shown in Figure 4B and 4C.

**Figure 4:**
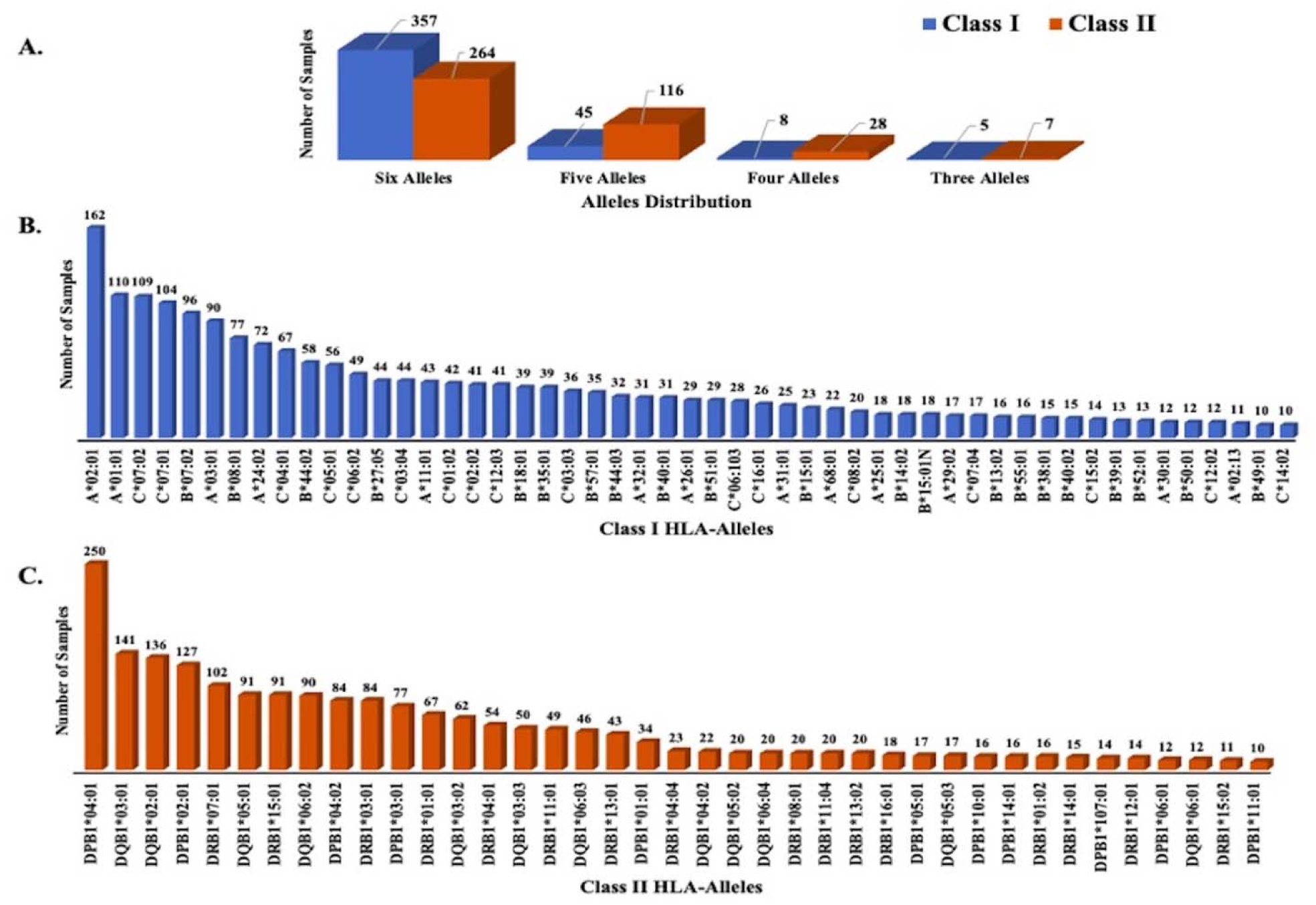
Frequency distribution of HLA-alleles in SKCM-samples, A) Describes the distribution of alleles in melanoma samples B) Describes the frequency of Class I HLA-allele with frequency >=10 C) Represents the frequency of Class II HLA -alleles with frequency >=10 samples

### 3.2 Categorization of Superalleles into Favourable and Unfavourable Groups

In order to understand whether an allele is favourable for survival of the patient or not, we compute difference in mean overall survival (MOS) of patients having and lacking a given allele (Table S3). Allele is assigned as favourable, if difference in MOS is positive, otherwise unfavourable. For instance, Class Allele HLA-A*01:01 is present in 110 patients with MOS 72.21 months; while, MOS is reduced to 55.25 months in 291 patients that lack it. It means this is a favourable allele as its presence enhance the MOS. Similarly, Class I allele HLA-A*24:02 present in 72 patients with MOS 43.73 and it is absent 329 patients with MOS 63 months. This is an unfavourable allele as its presence decreases the MOS of patients. There are several favourable and unfavourable alleles in both class of alleles as given in Supplementary Table S3. These alleles can be used to predict risk of survival, unfortunately, this statistics is biased as the number of patients having a particular allele is very small for most of the alleles. This prompted us to create the superalleles from these alleles. Therefore, HLA alleles were further assigned to superalleles on the basis of similarity in the HLA-genes and Field1 (F1). Here, 367 alleles were further categorized into 121 Superalleles. Out of 121 Superalleles, 60 and 61 belong to class I and II, respectively. HLA-A*01/02, HLA-B*07, HLA-C*07, HLA-B*44, HLA-DPB1*04/02, HLA-DQB1*02/03/06/05, HLA-DRB1*07/15 are the most frequent class I and class II HLA superalleles in the SKCM-patients represented in Supplementary Figure S1. Distribution of superalleles which are present in at least ten patients is shown in Figure 5. The abundance of all remaining superalleles is given in Supplementary Table S4.

**Figure 5:**
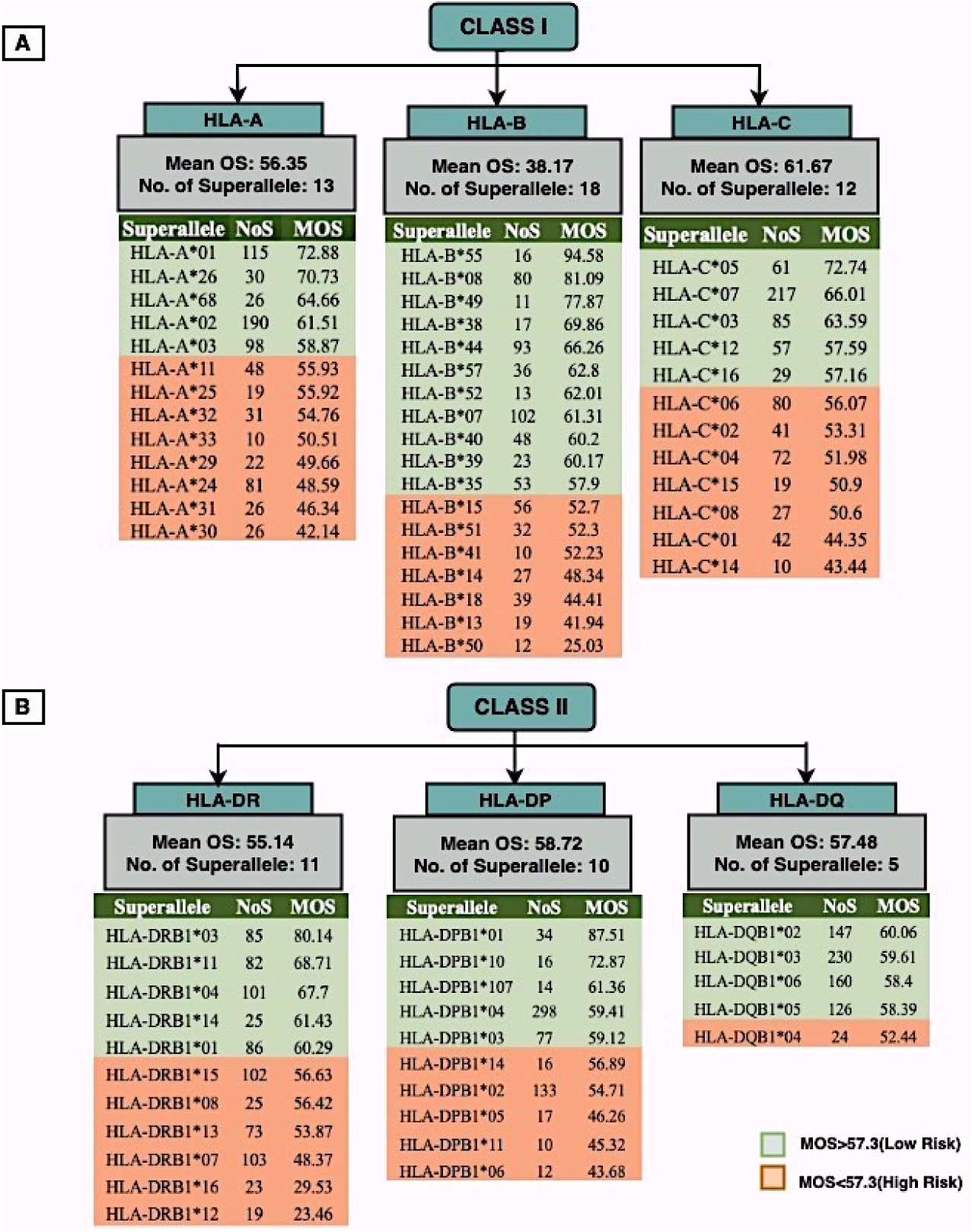
Distribution of HLA-superalleles in present in at least 10 SKCM-patients, A) Class I (B) Class II; MOS represents mean overall survival and NOS as number of samples/patients

The superalleles having MOS greater than 57.30 and lower than 57.30 are shown in Figure 5 with different colors. As shown in Figure 5, MOS of HLA-B superallele is the highest, i.e. 59.50 months among all other alleles in SKCM-patients, it means presence of this gene is favourable in OS of patients. Further, the HLA-superalleles are categorized into two groups, *i.e.* Survival Favourable (SF) and Survival Unfavourable (SU) on the basis of the difference in MOS between patients with a specific HLA-superallele-genotyping and patients lacking it. Among the 24 superalleles, 9 were SF (HLA-B*55, HLA-DPB1*01, HLA-DPB1*10, HLA-B*08, HLA-B*49, HLA-A*01, HLA-DRB1*03, HLA-C*05, HLA-C*07) and 15 were SU (HLA-B*14, HLA-A*24, HLA-DPB1*05, HLA-A*31, HLA-DPB1*11, HLA-DRB1*07, HLA-DPB1*06, HLA-C*14, HLA-B*18, HLA-C*01, HLA-B*13, HLA-A*30, HLA-DRB1*16, HLA-B*50, HLA-DRB1*12) with their mean overall survival and frequency are represented in Table 1.

**Table 1.** Classification of HLA-superalleles in to SF and SU on the basis of mean OS difference.

| HLA-superalles | #No. of Samples |  | #Mean OS |  | Mean Diff OS (P-A) | Class (Risk Status) |
| --- | --- | --- | --- | --- | --- | --- |
|  | Present | Absent | Present | Absent |  |  |
| HLA-B*55 | 16 | 385 | 94.58 | 58.46 | 36.12 | Favorable Super alleles (Low Risk) |
| HLA-DPB1*01 | 34 | 367 | 87.51 | 57.34 | 30.17 |  |
| HLA-B*08 | 80 | 321 | 81.09 | 54.62 | 26.47 |  |
| HLA-DRB1*03 | 85 | 316 | 80.14 | 54.46 | 25.69 |  |
| HLA-B*49 | 11 | 390 | 77.87 | 59.39 | 18.48 |  |
| HLA-A*01 | 115 | 286 | 72.88 | 54.68 | 18.20 |  |
| HLA-C*05 | 61 | 340 | 72.74 | 57.60 | 15.15 |  |
| HLA-DPB1*10 | 16 | 385 | 72.87 | 59.36 | 13.51 |  |
| HLA-C*07 | 217 | 184 | 66.01 | 52.70 | 13.31 |  |
| HLA-B*14 | 27 | 374 | 48.34 | 60.74 | -12.39 | Unfavorable Super alleles (High Risk) |
| HLA-A*24 | 81 | 320 | 48.59 | 62.77 | -14.18 |  |
| HLA-DPB1*05 | 17 | 384 | 46.26 | 60.51 | -14.25 |  |
| HLA-A*31 | 26 | 375 | 46.34 | 60.84 | -14.5 |  |
| HLA-DPB1*11 | 10 | 391 | 45.32 | 60.27 | -14.95 |  |
| HLA-DRB1*07 | 103 | 298 | 48.37 | 63.89 | -15.51 |  |
| HLA-DPB1*06 | 12 | 389 | 43.68 | 60.40 | -16.72 |  |
| HLA-C*14 | 10 | 391 | 43.44 | 60.32 | -16.88 |  |
| HLA-B*18 | 39 | 362 | 44.41 | 61.57 | -17.16 |  |
| HLA-C*01 | 42 | 359 | 44.35 | 61.72 | -17.37 |  |
| HLA-B*13 | 19 | 382 | 41.94 | 60.79 | -18.86 |  |
| HLA-A*30 | 26 | 375 | 42.14 | 61.13 | -19.00 |  |
| HLA-DRB1*16 | 23 | 378 | 29.53 | 61.75 | -32.22 |  |
| HLA-B*50 | 12 | 389 | 25.03 | 60.98 | -35.95 |  |
| HLA-DRB1*12 | 19 | 382 | 23.46 | 61.71 | -38.26 |  |

#Samples (P): No of SKCM-patients in which HLA-superallele is present; # Samples (A): No of SKCM-patients in which HLA-superallele is absent; #Mean OS (P): Average OS in which HLA-superallele is present ; # Mean OS (A): Average OS in which HLA-superallele is absent; Mean Diff OS: Mean difference in mean OS between patients with a specific HLA-superallele-genotyping and patients without it; Class: Survival Favourable (SF) or Survival Unfavourable (SU) HLA-superallele, SF considered as low-risk and SU taken as high-risk groups.

### 3.3 Univariate Survival Analysis

#### 3.3.1 HLA-Superalleles

It is clear from the above analysis that certain allele/superallele are responsible for improving the survival of patients. Next challenge is to utilize this information for predicting the high-risk cancer patients based on the presence of certain alleles or superalleles. Here, we used only superalleles for predicting the high-risk patients employing univariate survival analysis due to poor distribution of alleles in patients. We observed that HLA-B*50 which is responsible for poor survival of patients; assigned patients on high risk if this superallele is present and obtained HR 2.77 (95% CI 1.284 to 5.941) with p-value 0.009. Similarly, HLA-DRB1*12 achieved maximum performance HR 3.13 (95% CI 1.687 to 5.826) with p-value<0.001. The combined effect of the presence of HLA-B*50 and HLA-DRB1*12 is also used to predict high-risk patients and obtained HR 3.15, 95% (CI 1.906 to 5.194) with p-value less than 0.001, see Supplementary Table S5.

#### 3.3.2 Risk Score

To further improve the performance of the prediction models, we developed a threshold-based method using multiple superalleles as input features. In this case, we employed multiples variables that include both favourable and unfavourable superalleles. Towards this, first, we assign -1 and +1 for each favourable and unfavourable superallele, respectively. Thereafter, all the scores are cumulatively added to generate a single score called “Risk Score” for each patient. Further, to understand how well Risk Score based on superalleles stratified risk-groups of melanoma patients, survival analysis was performed using Risk Score as a input feature. For instance, if the threshold value is >=2 then the patients significantly divided into high-risk and low-risk groups with more than two-folds, i.e. HR 2.18 (95% CI 1.441 to 3.297) with p-value = 0.000223 as given in Table 2. Conclusively, we found that Risk Score thresholds act as a prognostic indicator for stratifying melanoma patients into high-risk and low-risk groups, as shown in Table 2. Additionally, Kaplan-Meier (KM) survival plots represent the segregation of high-risk and low-risk melanoma patients based on different threshold values of Risk Score, with significant p-values as shown in Figure 6.

**Table 2.** Survival analysis based on Risk score to discriminate low-risk and high-risk samples.

| Threshold<br>(Risk Score) | #G1 | #G2 | HR | 95% CI | P-value |
| --- | --- | --- | --- | --- | --- |
| $\geq 3$ | 375 | 26 | 1.84 | 0.966-3.508 | 0.0635 |
| $\geq 2$ | 341 | 60 | 2.18 | 1.441-3.297 | 0.000223*** |
| $\geq 1$ | 275 | 126 | 1.82 | 1.331-2.496 | 0.000183*** |
| $\geq 0$ | 171 | 230 | 1.71 | 1.277-2.302 | 0.000335*** |
| $\geq -1$ | 98 | 303 | 1.55 | 1.108-2.156 | 0.0103* |
| $\geq -2$ | 61 | 340 | 1.26 | 0.866-1.819 | 0.228 |
| $\geq -3$ | 37 | 364 | 1.55 | 0.977-2.463 | 0.06 |

#G1: No of SKCM-patients representing low-risk group; #G2: No of SKCM-patients denoting high-risk group; HR: Hazard Ratio; 95% CI: 95% confidence interval

**Figure 6.**
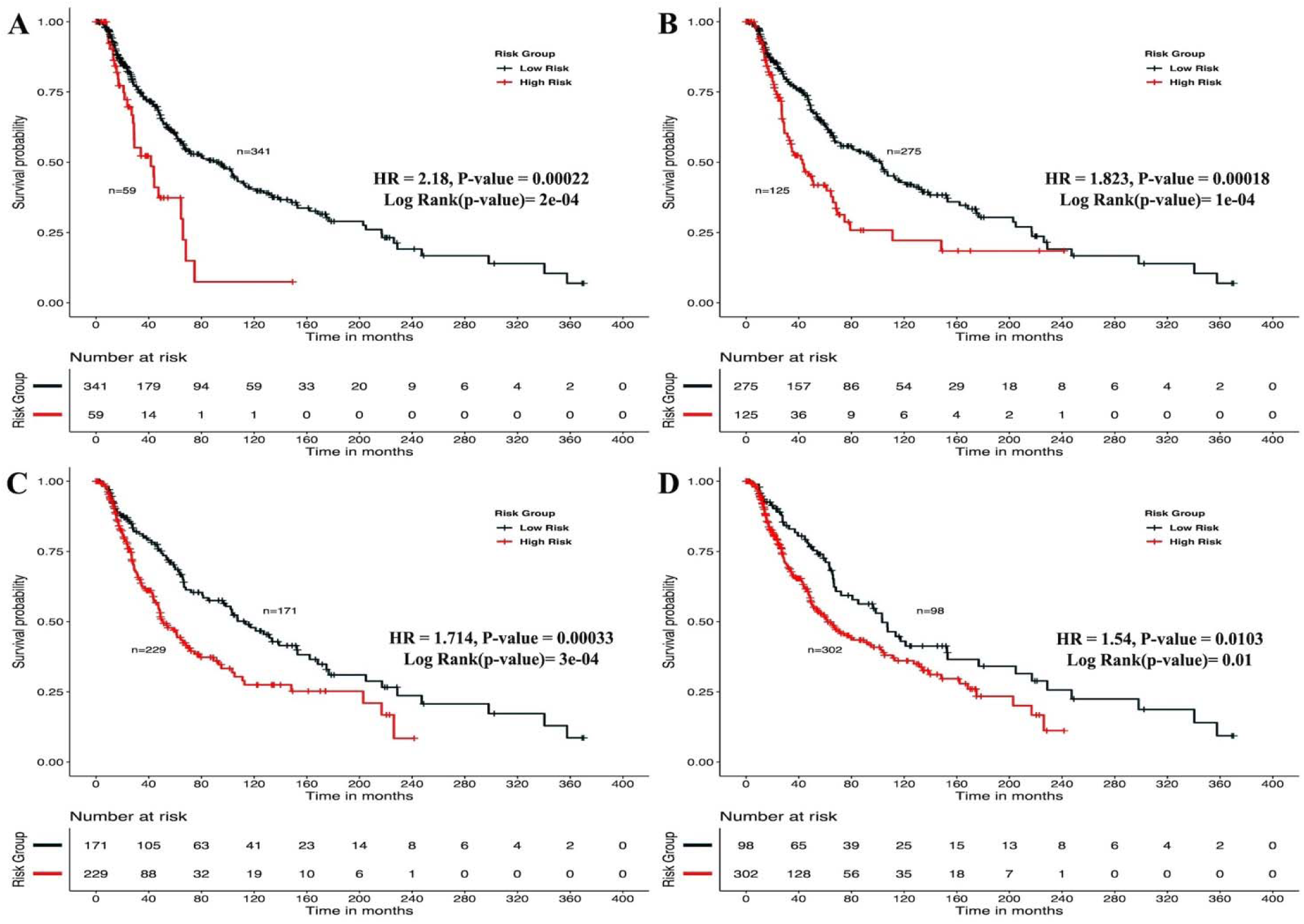
Kaplan Meier (KM) survival curves for the risk estimation of melanoma patient cohort based on the risk score with significant p-value (A) Melanoma samples stratified on the basis of cut-off (>=2 Risk Score), (B) Stratified samples by taking cut-off (>=1 Risk Score), (C) Stratified samples by taking cut-off (>= 0 Risk Score), (D) Stratified samples by taking cut-off (>=-1 Risk Score)

#### 3.3.3 Clinical Characteristics

In the past, the clinical features like age, gender, tumor stage, tumor status and Breslow depth have been shown a significant effect on the skin cancer incidence and bias towards a particular group (39). For instance, even in the current study, the male incidences are higher than of females in case of melanoma as shown in Supplementary Table S1. This prompted us to analyse the association between these clinical features and the survival of the patients. Thus, we perform the univariate survival analysis for the clinical features. This analysis indicates that the tumor status is one of the major significant prognostic factors in the prediction of survival of melanoma. Here also, we used threshold-based approach where we assign score +1 in case tumor is present in patient otherwise zero. We predict patient high-risk if score is more than zero and obtained HR 8.293 (95% CI 4.688-14.67) with p-value less than 0.0001 (Supplementary Table S6). Besides, age, tumor stage and Breslow depth are other clinical features that are significantly associated with the prognosis of the patients as shown in the KM plots shown in Figure 7. But, notably samples unable to stratified into high-risk and low-risk significantly based on the gender as represented in Figure7.

**Figure 7:**
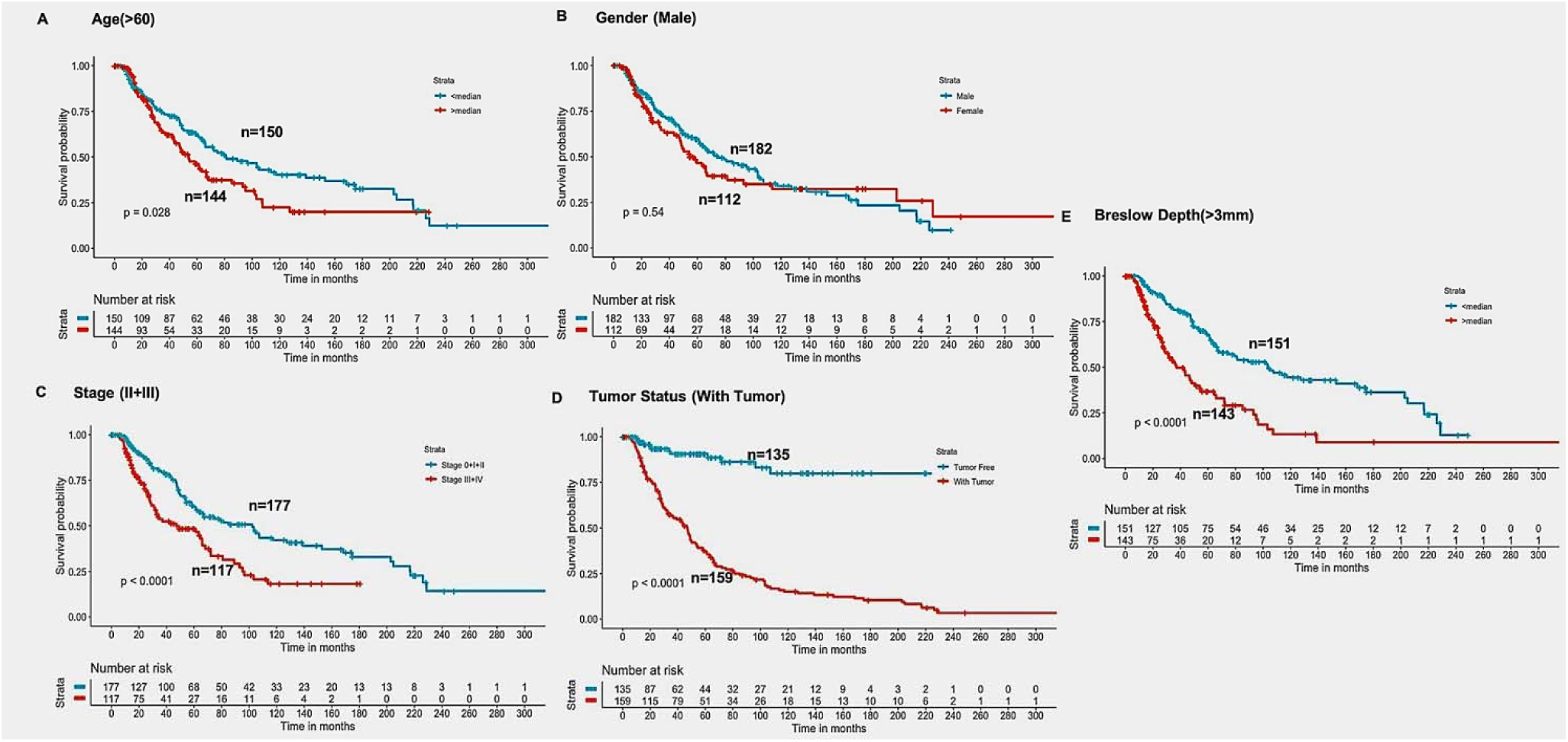
Kaplan Meier survival curves for risk estimation of SKCM cohort, show a significant difference in the high-risk/low-risk groups. (A)Patients with age (>60years) are stratified into high/low risk with HR=1.45, 95%CI=1.039-2.024 and p-value=0.028, (B) Stratification of low-risk and high-risk groups on the basis of gender with HR=1.11, 95%CI=0.7901-1.52, and p-value=0.545, (C) Stage (III+IV) patients are on high risk as compared to Stage (0+I+II) patients with HR=1.94, 95%CI=1.386-2.722, p-value<0.001, (D) Patients with Tumor status (With Tumor) were stratified on high/low-risk with HR=8.29, 95%CI=4.688-14.67, and p-value<0.001 (E) Patients having Breslow depth >3mm are stratified into high/low-risk corresponding 95%CI 1.788-3.509, HR=2.5, and p-value<0.001

### 3.4 Prediction Models

#### 3.4.1 Machine learning based prediction models

It is clear from the above results that both HLA-superalleles and clinical features (such as age, gender, tumor stage, tumor status and Breslow depth) are essential for the identification of high-risk patients. Though the threshold-based method is simple but not very efficient when we used multiple features. Thus, to further improve the performance, we implemented a wide range of machine learning techniques (e.g., Lasso, RF, Ridge, DT) for developing prediction models. The first model was developed by considering all factors including clinical as well as 24 HLA-superalleles. Lasso and RF based models obtained maximum performance with HR 3.17, p-value 3.50E-11, and HR 3.09, p-value 2.87E-11 for clinical features only, respectively, as shown in Table 3. Further, we developed models by eliminating two factors, i.e., tumor status and tumor stage, respectively. Since tumor stage is an important clinical factor, but this information is only available for a few patients. So, prediction model developed without considering these clinical factors, and achieved maximum HR=3.74 (with p-value=3.01E-14) by RF model. To further improve the performance of the machine learning based models, we used all clinical features and 24 HLA-superalleles. This model based on Lasso achieved maximum performance HR (4.05, 3.46, 3.51, and 3.11) with significant p-value for all four methods. Although, RF prediction models also performed reasonably well, but have lower HR than that of Lasso models. Complete results of the survival prediction models are represented in Table 3.

**Table 3.** Performance of the survival prediction models based on Clinical Characteristics and 24 HLA-Class I, II Superalleles implemented using various regression techniques.

| Method | Clinical Features Only |  | Clinical features + HLA-Superalles |  |
| --- | --- | --- | --- | --- |
|  | All Features |  |  |  |
|  | HR | P-value | HR | P-value |
| LASSO | 3.17 | 3.50E-11 | 4.05 | 4.01E-13 |
| RIDGE | 3.01 | 1.76E-10 | 3.80 | 2.30E-12 |
| RF | 3.09 | 2.87E-11 | 3.77 | 8.15E-12 |
| DT | 2.25 | 6.93E-07 | 2.00 | 5.29E-05 |
|  | Clinical Features without Tumor Status |  |  |  |
| LASSO | 3.50 | 3.93E-13 | 3.46 | 1.54E-11 |
| RIDGE | 3.49 | 3.93E-13 | 2.97 | 2.89E-09 |
| RF | 3.74 | 3.01E-14 | 2.96 | 8.23E-10 |
| DT | 2.15 | 2.24E-06 | 1.83 | 0.000312731 |
|  | Clinical Features without Tumor Stage |  |  |  |
| LASSO | 2.80 | 9.96E-10 | 3.51 | 1.32E-11 |
| RIDGE | 2.43 | 4.68E-08 | 3.55 | 7.56E-12 |
| RF | 2.81 | 2.05E-10 | 3.18 | 2.14E-10 |
| DT | 2.50 | 1.64E-08 | 2.76 | 2.38E-09 |
|  | Clinical Features without Tumor Stage & Tumor Status |  |  |  |
| LASSO | 2.40 | 4.41E-08 | 3.11 | 5.60E-10 |
| RIDGE | 2.40 | 4.41E-08 | 2.57 | 5.81E-08 |
| RF | 2.99 | 9.37E-12 | 2.59 | 1.55E-08 |
| DT | 2.54 | 1.06E-08 | 2.65 | 7.37E-09 |

HR: Hazard Ratio; RF: Random Forest; DT: Decision Tree

#### 3.4.2 Machine learning prediction models based on Wrapper method

It is important to have minimum number of features for avoiding over optimization and for the practical implementation in the real life. Therefore, further wrapper method used to decrease number of features recursively. In case of wrapper based recursive approach, one has to develop prediction model to evaluate performance after adding/removing a feature. Here, we used recursive machine learning method after addition of features approach. Finally, prediction models developed using five clinical features (age, gender, tumor stage, tumor status, and Breslow depth) and various HLA-superalleles by implementing different machine learning techniques (Table 4). Similar to above analysis, Lasso method based on five clinical features and 14 superalleles is the top performer with HR of 4.52 and p-value 8.01E-15 as given in Table 4. KM plot represents the stratification of high-risk and low-risk patients based on the estimated OS using Lasso recursive regression model as shown in Figure 8.

**Table 4.**
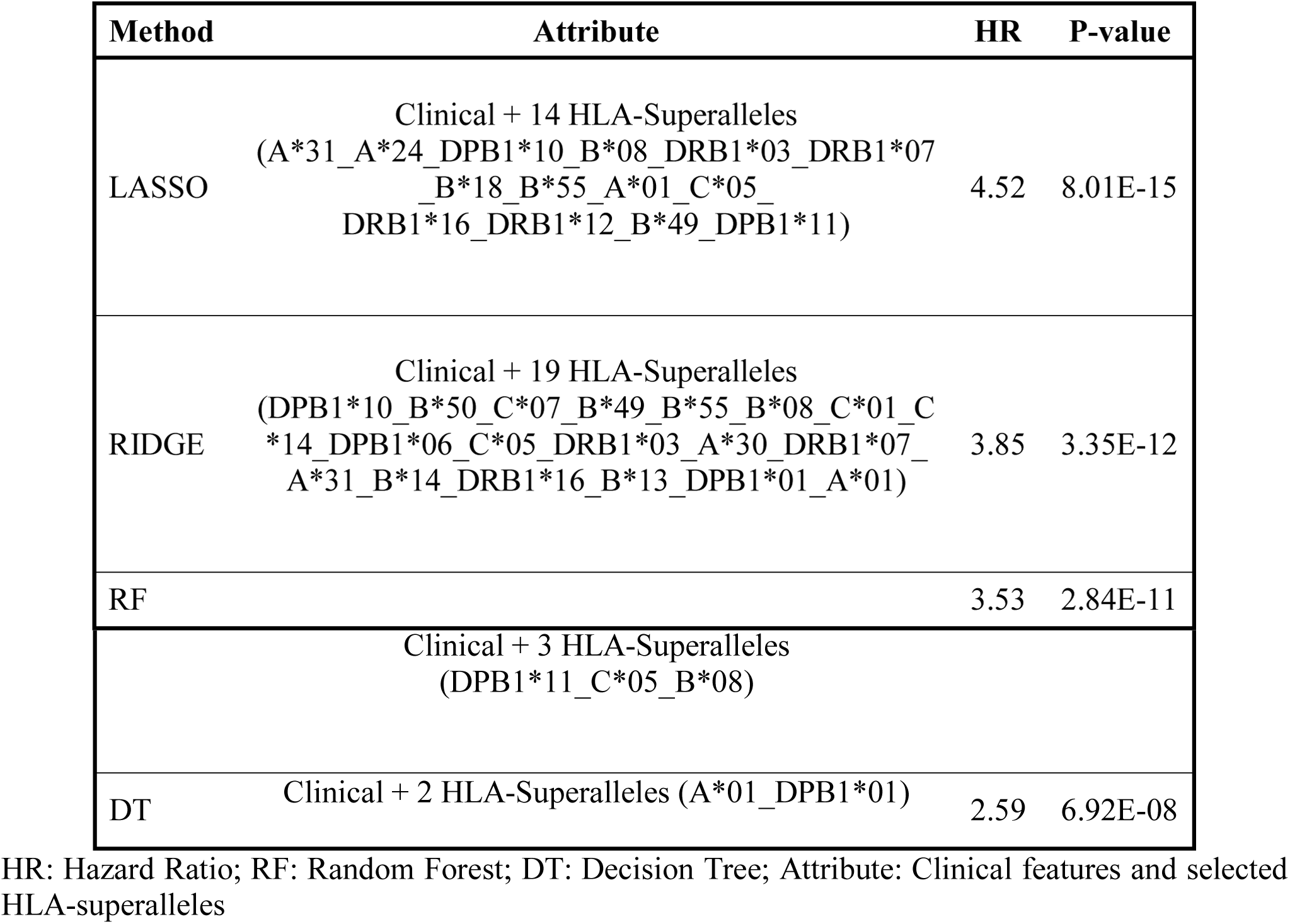
Performance of the recursive prediction models based on selected features (clinical features and superalleles) implemented using various regression technique.

**Figure 8:**
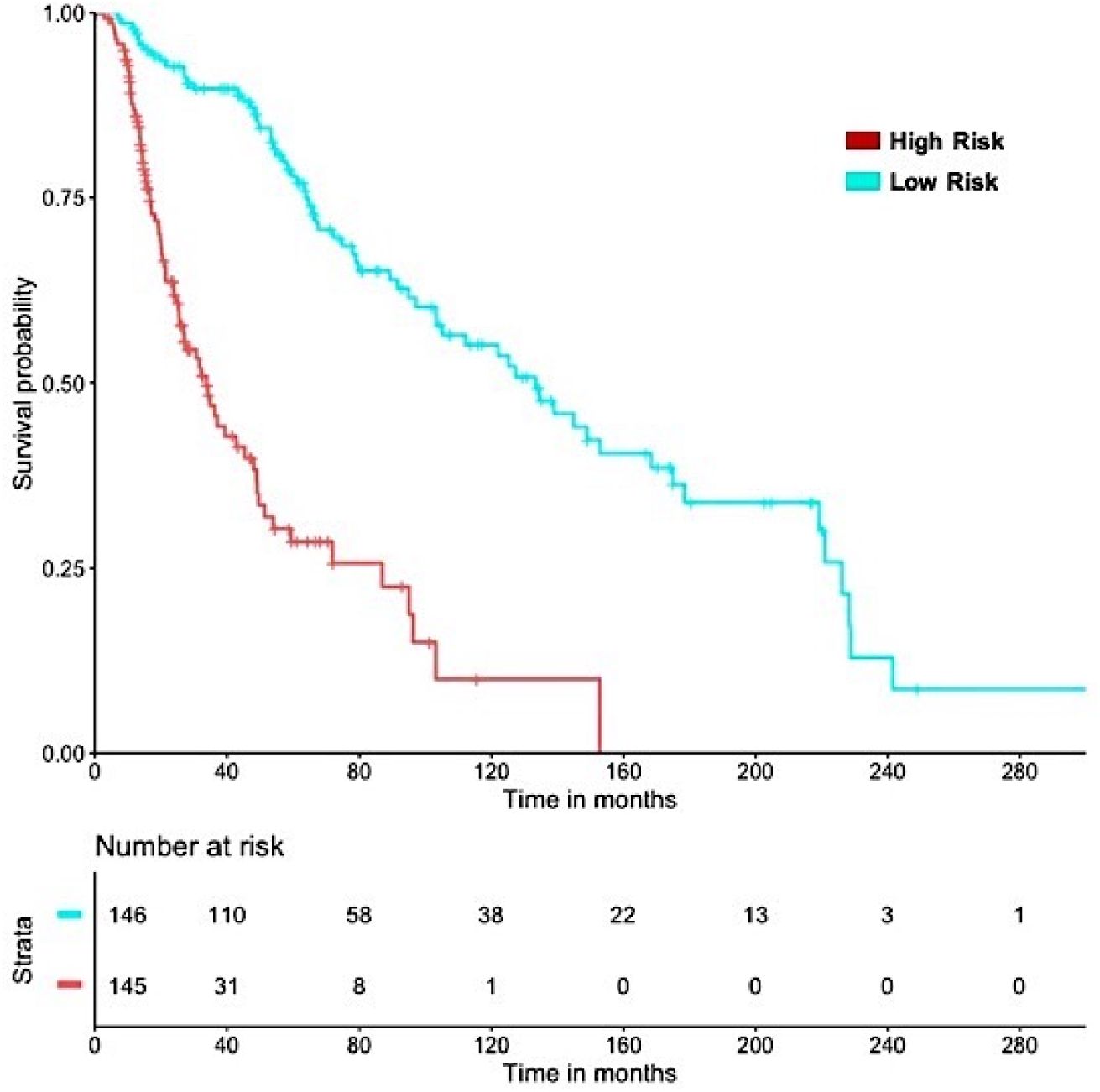
SKCM-patients were stratified based on predicted OS by using Lasso recursive regression model after applying five-fold cross validation. Samples with predicted OS < median (predicted OS) were at 4-fold higher risk as compared to the patients predicted OS > median (predicted OS) (HR = 4.52, 95% CI=3.088 to 6.609, p-value8.01E-15)

### 3.5 Multivariate Survival analysis for SF and SU HLA-superalleles

Further, to understand the combined effect of the multiple variables like SF and SU HLA-superalleles, Risk Score and clinical features on the survival of the patients, we perform multivariate survival analysis using Cox proportional hazard model (40). This analysis reveals that “Risk Score” is one of the most significant factors associated with the survival of patients. Results shown in Supplementary Figure S2, indicate that the presence of SU superalleles reduces the survival of melanoma samples. SU patients’ group is at approximately two times higher risk as compared to the SF patients’ group is indicated by HR = 2.44 (95% CI 1.68 to 3.5) with a p-value less than 3.02E-06, shown in Table 5. Both multivariate and univariate analysis reveals that age (>60), stage (III and IV), Breslow depth (>3mm) and Risk Score (>0) are associated with the poor survival in melanoma patients.

**Table 5:** Comparison of Univariate and Multivariate Analysis.

| Covariate | Univariate Survival Analysis |  |  | Multivariate Survival Analysis |  |  |
| --- | --- | --- | --- | --- | --- | --- |
|  | HR | 95% CI | P-value | HR | 95% CI | P-value |
| Age (>60years) | 1.45 | 1.039-2.024 | 0.029 | 1.45 | 1.03-2.0 | 0.032 |
| Gender(Female) | 1.11 | 0.790-1.520 | 0.545 | 0.98 | 0.69-1.4 | 0.896 |
| Tumor Stage(III+IV) | 1.94 | 1.386-2.722 | 0.001 | 1.89 | 1.33-2.7 | 0.0004 |
| Tumor Status(With Tumor) | 8.29 | 4.688-14.670 | <0.001 | 9.24 | 5.21-2.8 | 2.76E-14 |
| Breslow Depth(>3mm) | 2.50 | 1.788-3.509 | <0.001 | 1.96 | 1.38-2.8 | 0.00017 |
| Risk Score(>0) | 1.82 | 1.331-2.496 | <0.001 | 2.44 | 1.68-3.5 | 3.02E-06 |

HR: Hazard Ratio, 95% CI: 95% Confidence Interval

Further, to scrutinize which specific superalleles out of SF and SU superalleles groups, are significantly associated with good and poor outcome of the patients, multivariate analysis was performed using each of SF and SU superalleles with the clinical characteristics. Results from this analysis is shown that the presence of HLA-B*55 and HLA-A*01 superalleles is significantly associated with good outcome; while, HLA-DRB1*12, HLA-B*50, HLA-B*13, HLA-DPB1*06, HLA-A*31, HLA-A*24 are significantly associated with poor outcome of melanoma cohort in terms of their survival time as given in Supplementary (Table S7, S8 and Figure S3, S4).

### 3.6 Web Server for risk prediction in SKCM patients: SKCMhrp

To serve the scientific community, we developed a web server, “SKCMhrp” https://webs.iiitd.edu.in/raghava/skcmhrp/. SKCMhrp is designed for the risk prediction using clinical features and HLA-superalleles. It has two modules; one is based on clinical features and second is based on superalleles. First module predicts risk status of melanoma patients based on their clinical characteristics, *i.e.* age, gender, tumor stage, tumor status, Breslow depth. Here, an user can predict the survival time (in months) of the individual sample, even by choosing a single clinical feature. Input values are given to a regression model to estimate the Risk Status (RS). Second module predicts the risk status of melanoma patients using all 19 features that include five clinical characteristics and 14 superalleles.

## 4 Discussion

The American Cancer Society estimated 96,480 new melanoma cases (57,220 in men and 39,260 in women) in 2019, out of which around 7,230 people are expected to die (41). FDA has approved several therapies and strategies to curb melanoma over the past few years. Choosing a treatment from the available options requires information about the tumor such as its location, stage, *etc.* The major therapeutic options that exist are chemotherapy, radiotherapy, immunotherapy, photodynamic therapy, targeted therapy and surgical resection (42–44). Recent findings suggested that there is a relationship between the inadequate response of the immune system and the proliferation of melanoma cells (45). Antigenic repertoire variability is one of the crucial factors for the tumor progression and immunosurveillance (46). Due to the inadequate antigen processing mechanisms such as heterogeneous expression of HLA genes and defective immune system, it unable the CD8+ T-cells to recognize melanoma cells (47). HLA-Class I and II proteins are the key components of the immune system and have a significant role in the progression of melanoma(48,49). Gogas H et al., indicate that HLA-Cw*06-positive melanoma patients have better OS as compared to HLA-Cw*06-negative samples(50). Recent findings suggest that the higher expression of HLA-Class II genes enhances the survival of melanoma patients (51). This points out that the presence of HLA-Class II alleles also affects the survival of patients. Thus, it is important to understand which specific HLA-alleles from Class-I and Class-II could affect the survival of the patients. Eventually, how the HLA-genotypes of patients can improve the melanoma detection and therapeutic options for their better clinical outcome. The current study is a systematic attempt to understand the prognostic role of Class-I and Class-II alleles in the survival of melanoma patients. Towards this, firstly 367 HLA-genotypes identified for 415 skin cutaneous melanoma patients. These 367 alleles have lower frequency distribution among patients as shown in Table S1. Thus, it is difficult to delineate any reliable conclusion regarding any of the alleles from the analysis. This propels us for their assignment to 121 HLA-superalleles, based on the similarity of HLA-genes and Field1 (F1). Further, these superalleles are categorized into SF and SU groups based on the impact of their presence on the survival of the patients, *i.e.* higher MOS or lower MOS of the patients with their occurrence, respectively. Here, among the 24 superalleles, nine were SF include HLA-B*55, HLA-DPB1*01, HLA-DPB1*10, HLA-B*08, HLA-B*49, HLA-A*01, HLA-DRB1*03, HLA-C*05, HLA-C*07; while, 15 were SU that include HLA-B*14, HLA-A*24, HLA-DPB1*05, HLA-A*31, HLA-DPB1*11, HLA-DRB1*07, HLA-DPB1*06, HLA-C*14, HLA-B*18, HLA-C*01, HLA-B*13, HLA-A*30, HLA-DRB1*16, HLA-B*50, HLA-DRB1*12. Further, in the current study, Risk score is computed to evaluate the cumulative effect of the presence of SF and SU superalleles in patients. Thereafter, important HLA-superalleles, Risk score and clinical features like tumor status, age, and stage, were identified that can significantly stratified high-risk and low-risk survival groups employing univariate survival analysis and log rank test (Supplementary Table S3). Among them, particularly, HLA-B*50:01 (HR=2.77, p-value=0.01), HLA-DRB1*12:01 (HR=2.51, p-value=0.01), HLA-DPB1*05:02 (HR=2.39, p-value=0.01), HLA-C*15:02 (HR=1.91, p-value=0.05), HLA-B*35:01(HR=1.52, p-value=0.06) significantly reduces the OS.

In the current study, multivariate analysis reveals SF and SU HLA-superalleles as the independent prognostic indicators. For instance, the presence of HLA-Class I superalleles include HLA-B*55 (HR=0.15, p-value=0.013) and HLA-A*01(HR=0.54, p-value=0.011)) are significantly associated with good outcome (Supplementary Table S7, Figure S4). On the other hand, superalleles such as HLA-B*50 (HR-3.35, p-value=0.02), HLA-DRB1*12 (HR=3.44, p-value=0.028), HLA-DRB1*16 (HR=2.18, p-value=0.04), HLA-B*13 (HR=2.49, p-value=0.04), HLA-DPB1*06 (HR=3.15, p-value=0.006), HLA-A*31 (HR=2.09, p-value=0.01) and, HLA-A*24 (HR=1.76, p-value=0.006) associated with the poor survival outcome in SKCM-cohort (Supplementary Table S8, Figure S5). Eventually, the multivariate analysis revealed the Risk score, tumor status, tumor stage, Breslow depth and age as major independent prognostic factors for melanoma patients. Besides, the low expression (with mean cut-off) of HLA (-A, -B, -C, -DPB1, -DQB1, -DRB1) genes, consequently decreases the OS rate of melanoma cohort shown in Supplementary Table S9. Furthermore, various prediction models developed for the estimation of survival time of patients based on clinical characteristics, HLA-superalleles genotypes, and their various combinations implementing different machine learning techniques, *i.e.* Lasso, Random forest, DT and Ridge regression models. Subsequently, the predicted OS from these machine learning algorithms further employed for the stratification of high-risk and low-risk survival groups. Although, the prediction based on five clinical factors attained consistent performance, i.e. HR=3.17; but, stage and tumor status are two important factors which are mostly not available as their determination is a difficult task. Therefore, we have also developed prediction models after exclusion of these two factors. The performance of our ML models substantially decreased to HR 2.40. Further, prediction models developed employing important clinical factors with HLA-superalleles and removing tumor stage and tumor status as well. Results indicate that the performance of models based on HLA-superalleles and conveniently available clinical factors like age, gender and Breslow depth considerably improved from HR (2.40 to 3.11). Lasso and RF recursive regression models are among the top performers to predict survival of melanoma samples. Particularly, predicted OS obtained from Lasso recursive model, based on clinical characteristics and nine-superalleles significantly (p-value<0.001) stratified the high-risk and low-risk survival groups of the melanoma patients with HR=4.52. Although, RF-based models performed reasonably well in the estimation of OS, but, stratified survival risk groups with lower performance than that of Lasso models, *i.e.* HR=3.53 only.

## 5 Conclusion

Taken together, our findings exhibit that the occurrence of HLA-Class I, II alleles genotype influence the overall survival of SKCM patients both in favourable and unfavourable directions. Eventually, survival analysis and recursive machine learning regression models revealed the prognostic potential of important 14 superalleles and five clinical factors in the stratification of high-risk and low-risk survival groups and the estimation of overall survival time, respectively. Further, these HLA-based signatures could be considered to design personalized vaccine in several clinical cohorts. For the clinical utility, this further needs to confirm by exploring the role of these superalleles in other cohorts. Finally, to provide service to scientific community for prediction of high-risk patients based on their clinical features and 14 HLA-superalleles, we designated webserver “SKCMhrp”.

## 7 Author Contributions

AD, HK, SB, collected the data and processed the datasets. AD, HK, SP, implement the algorithm. AD, SP, created the back-end server and front-end user interface. AD, SP, developed prediction models. AD, HK and GPSR analysed the results. AD, HK, SP, CA, and GPSR penned the manuscript. GPSR conceived and coordinated the project, facilitated in the interpretation and data analysis and gave overall supervision to the project. All authors have read and approved the final manuscript.

## 9 Data Availability Statement

All the datasets generated for this study are either included in this article/Supplementary material or available at the “SKCMhrp” webserver https://webs.iiitd.edu.in/raghava/skcmhrp/data.php, as mentioned in the Materials and Methods section.

## 10 Acknowledgements

All the authors acknowledge funding agencies J. C. Bose National Fellowship (DST). AD, SP, HK, and SB, are thankful to DST INSPIRE, DBT, CSIR and ICMR for providing fellowships, respectively.

## 11 Supplementary Material

The supplementary material for this article can be found at………

