## Supplementary Figure S1, S2, S3, S4 for "Computing Skin Cutaneous Melanoma Outcome from the HLA-alleles and Clinical Characteristics"

**Supplementary Figures**

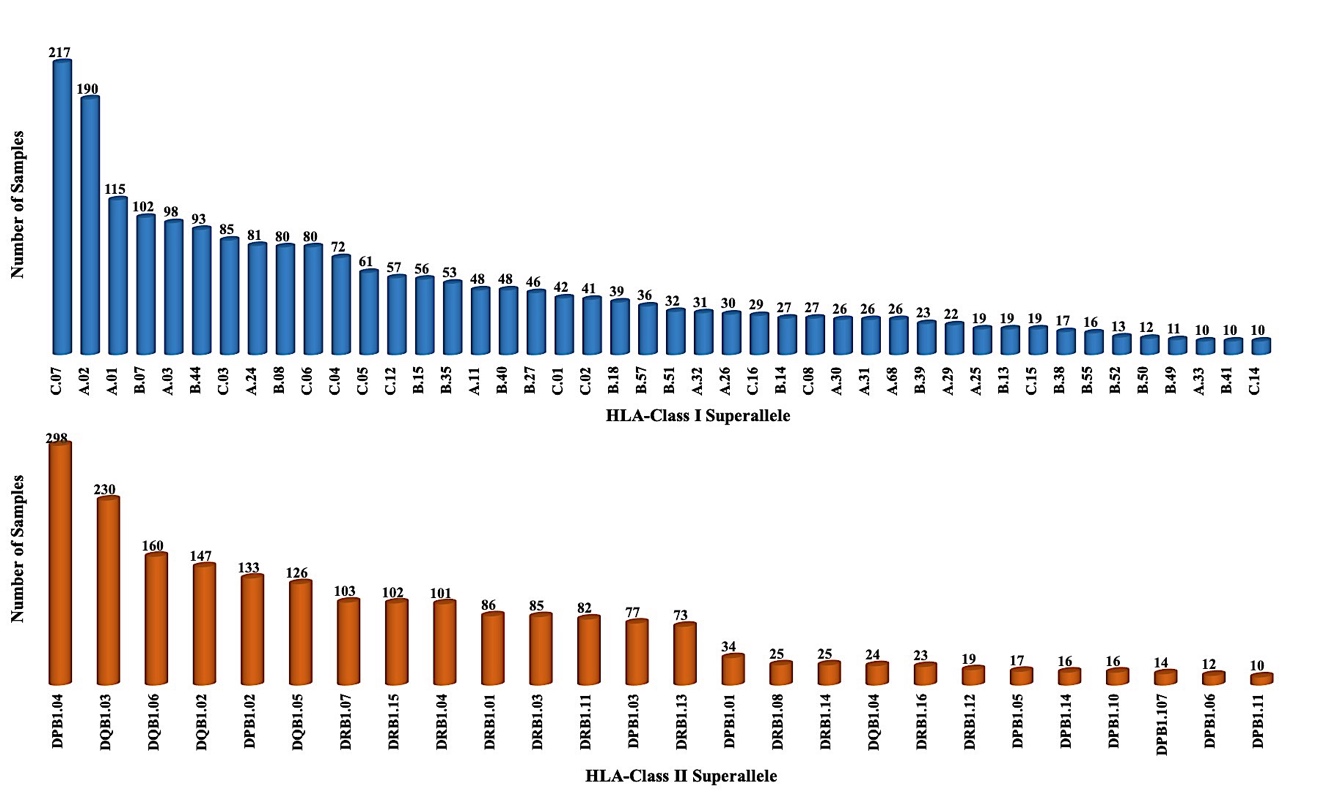
**Figure S1** Abundance of Class I and II HLA-Superalleles in SKCM cohort with a frequency more than or equal to 10 patients

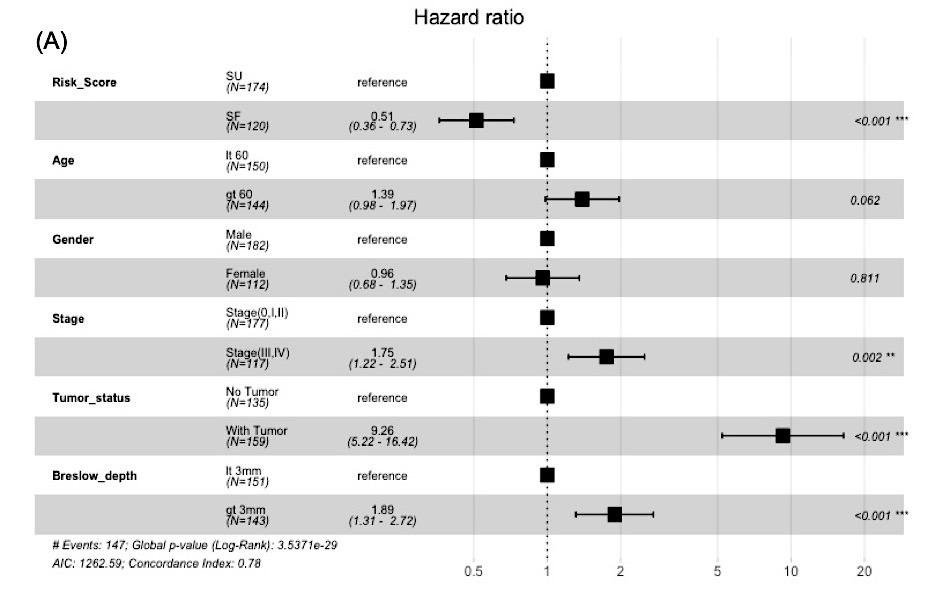

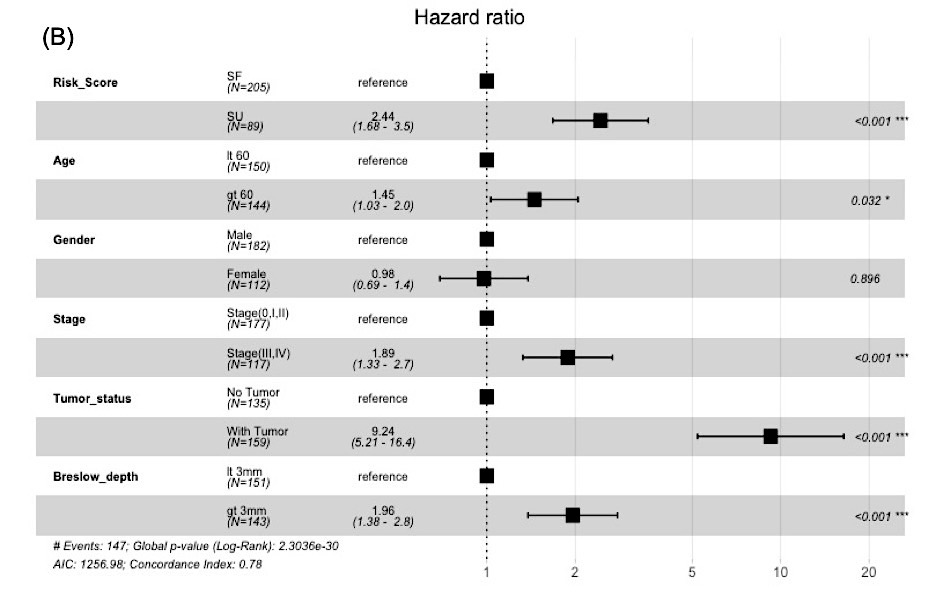

**Figure S2** Multivariate Cox-PH hazard model for risk estimation in SKCM-patients based on (A) presence of SF superalleles (Risk Score >0) with HR=0.51(0.36-0.73), p-value<0.001 and (B) presence of SU superalleles (Risk Score <0) cut-offs with HR=2.44(1.68-3.5), p-value<0.001.

**Figure S3** Multivariate Cox-PH hazard model for risk estimation in SKCM-patients based on presence of SF superalleles and clinical features. Presence of superalleles (HLA-B*55 and HLA-A*01) significantly increases the overall survival of patients
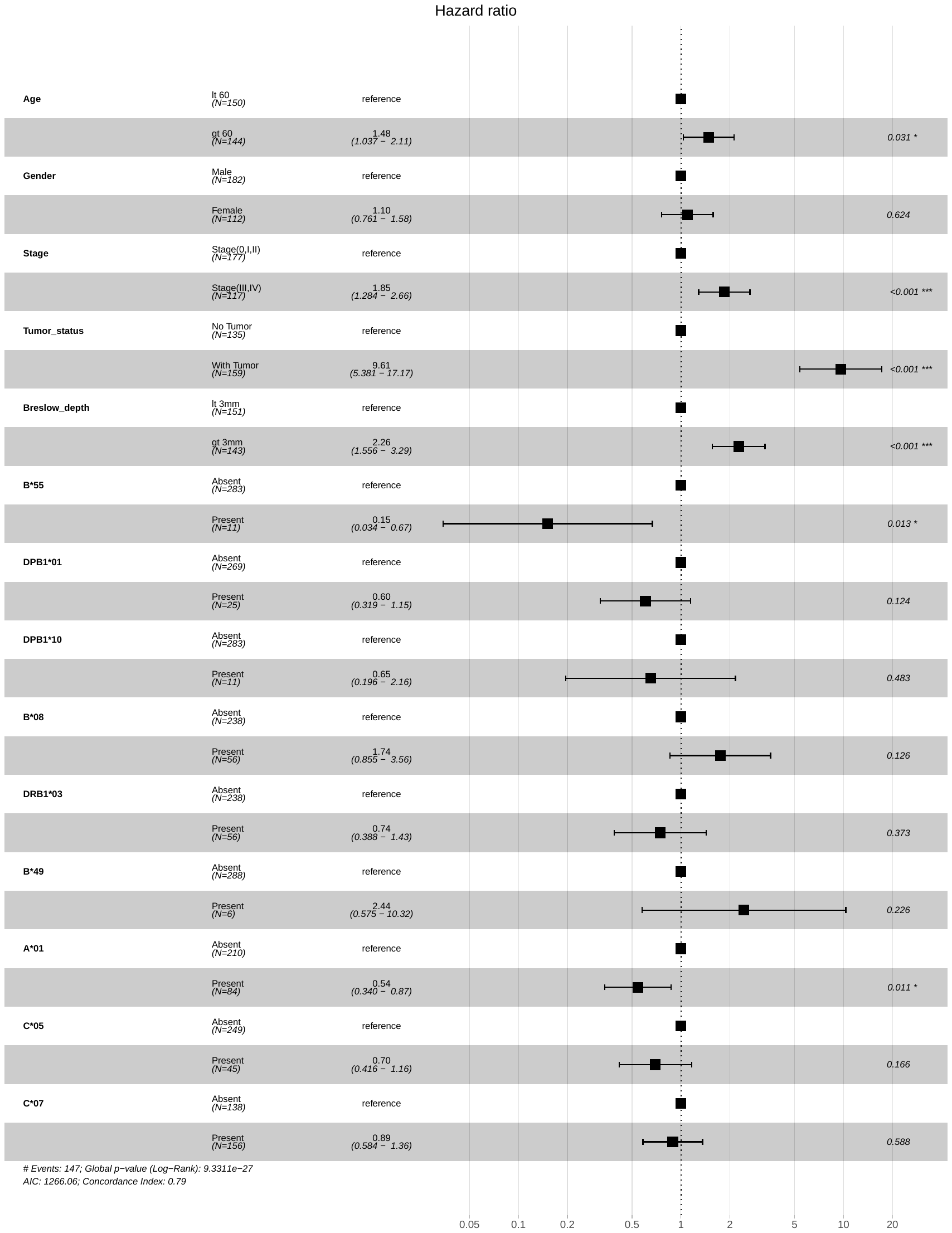
.

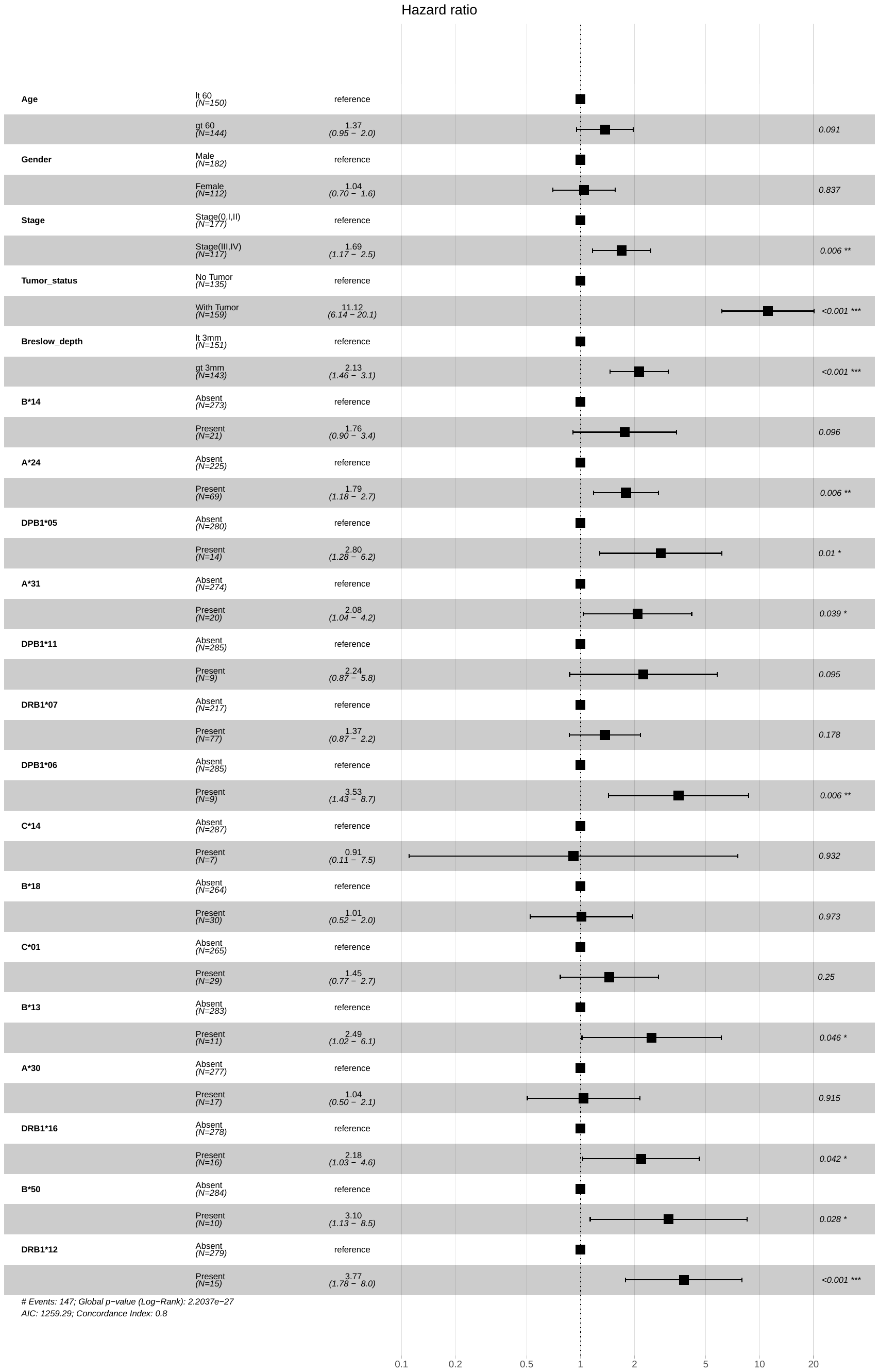

**Figure S4** Multivariate Cox-PH hazard model for risk estimation in SKCM-patients based on presence of SF superalleles and clinical features. Presence of superalleles (HLA-DRB1*12 (HR=3.77, p-val<0.001), HLA-B*50 (3.10,p-val), HLA-B*13 (HR=2.49,p-val=0.04), HLA-DPB1*06 (HR=3.53, p-val=0.06), HLA-A*31(HR=2.08,p-val=0.032), HLA-A*24 (1.79,p-val=0.006)) significantly stratifies the high/low risk groups.
